# A preserved neural code for temporal order between memory formation and recall in the human medial temporal lobe

**DOI:** 10.1101/2024.10.12.618011

**Authors:** Mohammad Farhan Khazali, Armin Brandt, Peter C. Reinacher, Michael Kahana, Joshua Jacobs, Andreas Schulze-Bonhage, Lukas Kunz

**Affiliations:** Epilepsy Center, Medical Center – University of Freiburg, Faculty of Medicine, University of Freiburg, Freiburg, Germany; Department of Stereotactic and Functional Neurosurgery, Medical Center – University of Freiburg, Faculty of Medicine, University of Freiburg, Freiburg, Germany; Fraunhofer Institute for Laser Technology, Aachen, Germany; Department of Psychology, University of Pennsylvania, Philadelphia, PA, USA; Department of Biomedical Engineering, Columbia University, New York, NY, USA; Department of Epileptology, University Hospital Bonn, Bonn, Germany

## Abstract

Temporal memory enables us to remember the temporal order of events happening in our life. The human medial temporal lobe (MTL) appears to contain neural representations supporting temporal memory formation, but the cellular mechanisms that preserve temporal order information for recall are largely unknown. Here, we examined whether human MTL neuronal activity represents the temporal position of events during memory formation and recall, using invasive single and multi-unit recordings in human epilepsy patients (n = 19). Participants freely navigated a virtual environment in order to explore and remember locations and temporal positions of objects. During each exploration period, they sequentially encountered two or three different objects, placed in different locations. This allowed us to examine single- and multi-unit neuronal firing rates (FR) as a function of the temporal position the objects were presented in. We found that a significant number of multi-units and single-units in various MTL regions including the hippocampus showed selectivity to the temporal position of objects during the exploration period. During recall, patients were asked to indicate which one of two objects from the same trial was found latter. Neural firing rates during recall showed a selectivity supporting recall of temporal positions. Interestingly, most of the selective single-units that stayed selective during encoding and recall preserved their temporal position preference. Our results thus suggest that neuronal activity in the human MTL contains a preserved neural code for temporal order in memory formation and recall.

## Introduction

Our lives are organized in time, therefore events happening in our lives follow a temporal order. Thus, when the brain form memory traces about events in our lives, they aim to maintain the same temporal order because it is ecologically beneficial. Amnesia for temporal order and content were observed in patients with thalamus lesions (*1*, *2*). Furthermore, bilateral damage to the fornix, eliminating communication between frontal lobe and hippocampus, was found to cause retrograde temporal order amnesia without affecting non-temporal memories (*4*). The medial prefrontal and retrosplenial cortices have also been reported to play a role in recency memory in rodents (*4–6*). In the context of working memory, the order of items in a sequence is encoded by neural firing phase with respect to theta oscillations (*7*, *8*). In the context of memory that covers longer periods of time combined with other behavioral tasks such as navigation, previous human single-neuron studies in epilepsy patients observed neurons in the human medial temporal lobe (MTL) regions that track the time and temporal order of events (*9–12*). Together, these different lines of research suggest that the human brain contains dedicated neural mechanisms for temporal order memory.

Here, we aimed at understanding whether neural processes for temporal order during memory formation are related to those during retrieval. Specifically, it currently remains elusive whether neuronal activations that encode temporal order during memory formation get reactivated during temporal order recall. This would suggest a preserved neural code in a population of neurons that is involved in both computations of encoding and recalling. A prior study indicates that time cells, which are neurons that fire consistently in specific moment of time in a task, may be unrelated to this process as time cells were found to comprise two different populations that were active either during either the encoding or recall period of episodic memory (*13*). Though time cells’ activity during encoding predicted the temporal organization of retrieved memory, the activity of the encoding and recalling populations of time cells were segregated: Most of the time cells which were selective to time during encoding were not selective to time during recall, and a small proportion of the encoding population that stayed selective to time during recall did not show consistent activation similar to the one during encoding (*13*). Thus, how the encoded information about temporal order is transferred to neuronal populations supporting recall remains unknown.

Hence, in this study, we examined neuronal activity in the human MTL related to temporal memory encoding and recall in a virtual-reality task combining elements of navigation and episodic memory which required participants to recall the temporal order of items presented in a sequence. We investigated whether the transfer of temporal order information from encoding to recall might engage neuronal populations that are engaged in both processes. To maintain the temporal order information across both processes, we furthermore hypothesized that the selectivity exhibited by MTL neurons during encoding would be preserved during recall. Our results revealed three neuronal populations that are engaged in temporal order memory. The first group of neurons appeared to be involved in encoding temporal order, the second in recalling temporal order information, and the in both encoding and recalling temporal order information, predominantly at identical temporal positions. We finally show that a small number of neurons is enough to preserve the temporal code, potentially playing a central role in transferring the temporal order code between both processes.

## Results

To test our hypotheses, we used invasive neuronal recordings in human epilepsy patients. We recorded neural activity while the participating patients freely navigated a virtual environment to collect different objects hidden in treasure chests (so-called Treasure Hunt task, TH). The task required participants to encode and remember the identity of the objects, their locations, and their temporal positions.

During each trial, participants sequentially encountered two or three different objects, randomly placed in different locations (**Fig. 1A, B**). The identity of the objects changed on a trial by trial basis. After collecting these objects, participants either had to report the locations of where they collected the objects or to recall the object’s name if the location of where they were collected was indicated (**Fig. 1C**). Afterwards, they were asked to report the relative temporal order of the objects by selecting one of two objects they had collected later than the other during the same trial (**Fig. 1D**). We refer to the time needed to collect an object, including navigating toward and collecting it, as *epoch* (see **Fig. 1E**, for patient 12). On average, patients needed around 8.3 ± 2.8 seconds (mean ± standard deviation (std)) for each epoch. All patients showed a tendency for faster trial completion over time, indicated by a negative slope of a linear regression fit of the epoch duration to the trial index (**Fig. 1F**). Thus, participants became better in navigating the virtual environment with time. During the retrieval period, participants reported the temporal order of objects correctly with an average accuracy of 90 ± 9% (mean ± 95% confidence interval (ci)). Thus, our recordings from the MTLs of these patients offer a window into the neuronal mechanisms governing temporal memory encoding and recall.

**Figure 1.**
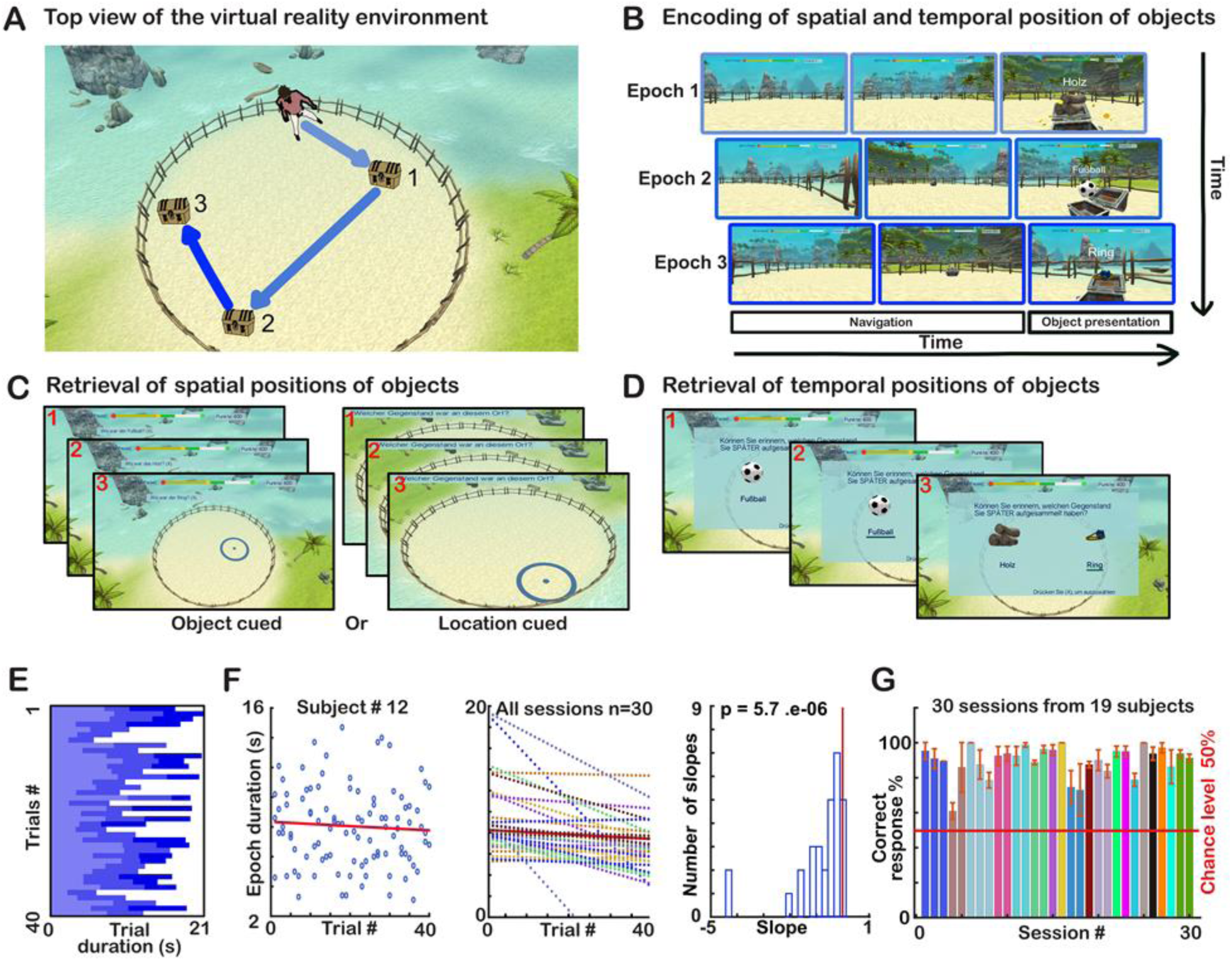
Behavioral evaluation of patients’ performance in the Treasure Hunt (TH) paradigm. **A**. Top view of the virtual reality in which the subjects had to navigate and collect two or three objects per trial, hidden in treasure chests that are presented in a temporal sequence (memory encoding). **B**. Subject view while navigating and collecting treasures. When navigation starts, epoch1 starts and the subject navigates the way until collecting treasure1. Then epoch2 starts until the collection of treasure2. That will indicate the start of epoch3 or the end of the whole navigation part (if the trial has just two treasures). **C**. Presents the spatial memory retrieval part of the task that is consisting of either: 1-asking the subject to indicate the location in which a specific treasure (object) was collected at (object cued trial) or by asking the subject to say the name of the object that was collected at a shown location (location cued trial). **D**. Presents temporal memory retrieval part, in which the subject is presented by two collected objects from the current trial and has to indicate the object collected latter. **E**. Shows the durations of each epoch per trial conducted in one session by subject 12. Note that epoch1 is indicated in very light blue, epoch2 in light blue and epoch3 in dark blue. **F.** Presents the relationship between epoch duration, time needed to collect an object in a trial and the trial index. **Left**, shows a tendency for less time needed to collect objects in later. **Middle** and **right**, on average most sessions showed that subjects needed less time to collect objects in later trials. **G.** Presents the accuracy percentage of correct retrieval responses of temporal positions, all sessions of a given patient are indicated in the same color. The response accuracy of all subjects was higher than chance level.

To carefully explore these mechanisms, we distinguished between single- and multi-unit neuronal recordings (SUI) and (MUI), based on the variability of their spike waveforms (**see methods**). To assess the quality of this classification, we examined the hypothesis that SUI recordings have a higher signal-to-noise ratio, because they putatively originate from a single neuron with consistent biophysical properties, in contrast to MUI recordings. Exploring three biophysical criteria suggested that our classification as SUI or MUI was accurate: First, higher peak-peak amplitude and second lower spikes counts for SUI (**Fig. S1A, B**). Peak-peak amplitude for SUI was around 107.1 ±39.3 µV (mean ± std) and 51.1 ± 10.4 µV for MUI (p<0.001; Wilcoxon rank sum test, **Fig. S1Ci**). Median spike count for SUI was 2495 (25%-75% ci: 945.5-7155.5) spikes and 8004 (3117–16978) spikes for MUI (p<0.0001; Wilcoxon rank sum test, **Fig. S1Cii**). The variability of the spike waveform was much lower for SUI than for MUI recordings. A linear regression fit of the spike waveform variability, measured by the standard deviation, to the peak-peak amplitude as input resulted in much steeper slope for MUI recordings as compared to SUI, (**Fig. S1Ciii)**. We confirmed this statistically by repeating the fitting process 100 times for 100 randomly chosen data points, bootstrapping with replacement. The resulting slopes for MUI data was significantly steeper with 0.1153 ± 0.022 (mean ± std) as compared to the ones of SUI 0.023 ± 0.004 (p<0.0001; Wilcoxon rank sum test).

After establishing the quantitative difference between both signals, we separately examined their relation to temporal memory encoding. We evaluated the SUI and MUI FRs as a function of the temporal position which the objects were presented in during the navigation period. Both signals exhibited significant preferences to different temporal order positions (**Fig. 2**). For example, a hippocampal single-unit recording (**Fig. 2Ai**) showed a clearly higher FR in epoch 1 (i.e., the first temporal position) during which the participant was required to navigate toward and collect the first object. The average FR of this neuron was 2.4 ± 0.6, 1.1 ± 0.4 and 0.9 ± 0.5 Hz (mean ± 95% confidence interval (ci)) for the first, second and third positions respectively with a significant different between the three positions (p<0.001; ANOVA-surrogate statistics, see methods).

**Figure 2.**
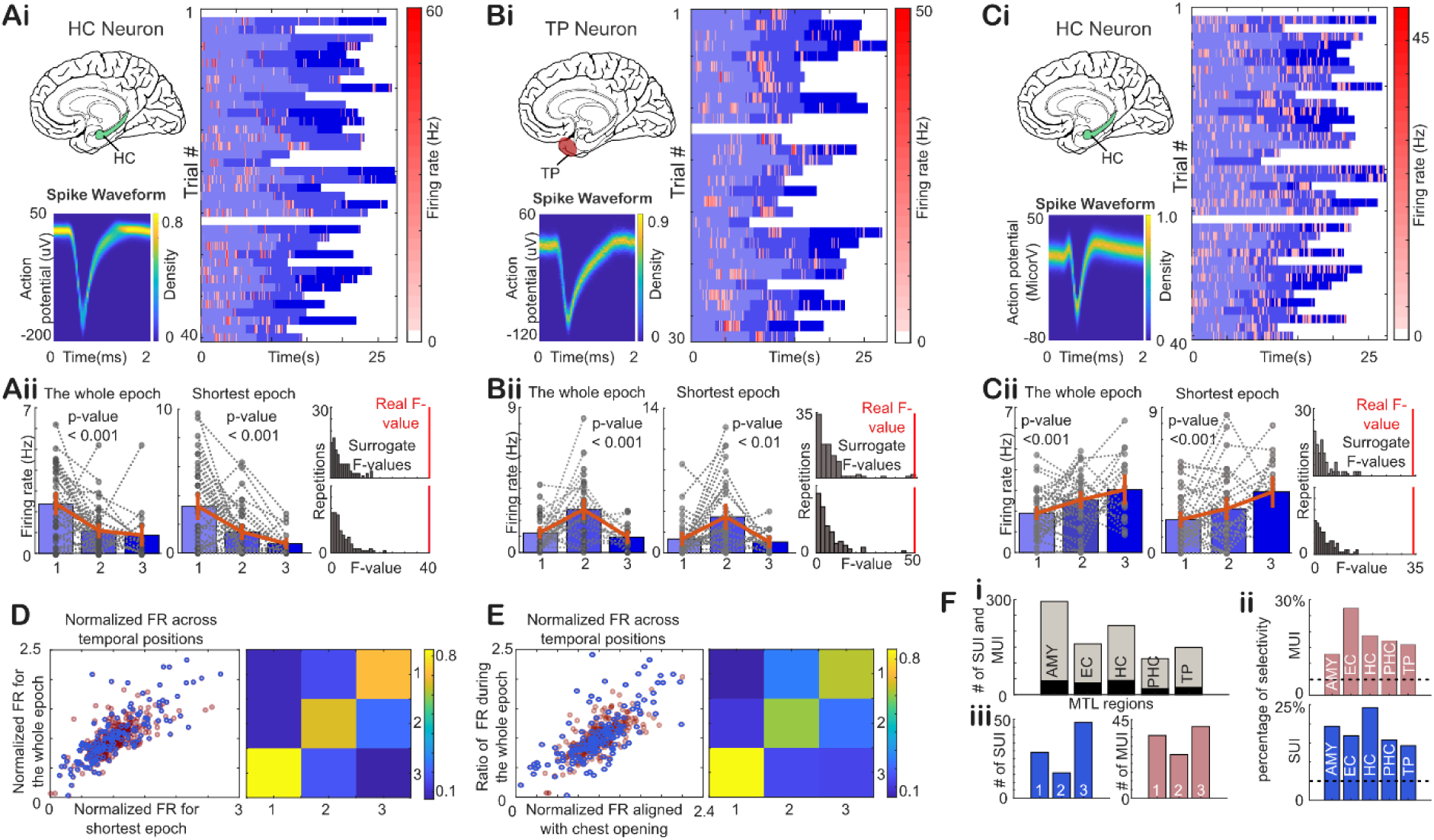
Neurons in the MTL are selective to different temporal positions during navigation. **Ai** Plots the spiking events within 100ms bins of a HC neuron, superimposed on the temporal positions (epochs) in blue (very light, light and dark blue for epoch 1, 2, and 3, respectively). Note that spiking events of the neuron are concentrated in epoch 1 and the ones of neuron B and C are concentrated in epochs 2 and 3, respectively. Note the density function plot in Ai which plots the spike waveform **Aii**. Quantifies the selectivity of the neurons. The left bar plot present the averaged firing rate during each epoch (colors indicate the epoch as in Ai. Black circles indicate the average firing rate per epoch per trial, black lines connect between them for the same trial. The right bar plot presents the averaged firing rate during each epoch showing the time needed for the fastest reaction time (RT). The two histogram plots at the right present raw F-value in red with respect to surrogate F-values generated from one-way ANOVA testing of shuffled data across the three epochs. The upper plot is generated for the shortest time needed for fastest RT. The lower plot is generated using the whole epoch. Note that both plots show similar and strong selectivity to the first epoch. **B** and **C** replicate the plots in A for two neurons preferring the second and third epoch, respectively. **D**. Plots the averaged firing rate for each epoch, normalized by the average firing rate across all epochs. The x-axis plots FRs averaged over the time of the shortest epoch against FRs averaged across the whole epoch on the y-axis. Thus each neuron provides three data-points, one for each epoch. Note the significant correlation between the FRs across both periods of time. The highest FRs across the epoch stay the same independently of the period chosen to average the FR, as indicated by the heat plot. **E**. Replicates D, but for FRs averaged across the whole epoch with FRs averaged from 3 seconds before chest open to 1.5 after chest open. **F**. Quantifies the overall neural recordings. **i**. Illustrates the number of MUIs and SUIs recorded from each region in gray and the number of selective UIs in black. **ii**. Plots the percentages of selective SUIs per region in pink and for MUIs per region in blue. **iii**. Plots the preferred epoch of each SUI in pink and MUI in blue. Note that both signals exhibit similar preferences to the first and third position.

Since the patients knew in which temporal position they were at, right from the beginning of each epoch, we hypothesized that temporal order information must be available already early during the epochs. To probe this hypothesis, we restricted our analysis of temporal order to the shortest time needed to finish any epoch, which we refer to as the *shortest epoch duration* (SED). For this neuron example, SED was 2.6 seconds. Hence, when averaging FR during the first 2.6 seconds of each epoch, despite the average length of all epochs being 8.2 ±33.9 seconds (mean ± std) for this session, the preferred temporal position stayed the same. Specifically, the FRs during SED were 3.2 ± 0.9, 1.5 ± 0.6 and 0.7 ± 0.4 (mean ± ci) for the first, second and third epoch, respectively, and the firing rates were significantly different from each other (p<0.001; ANOVA-surrogate statistics, see methods). Thus, the temporal position preference seems to be present already early during each epoch (**Fig. 2Aii**). This was not only the case for neurons preferring the first temporal position, but also for neurons preferring later positions (**Fig. 2B, C**).

We found that the temporal position selectivity was not limited to SUIs, but MUIs showed similar selectivity to different temporal positions (**Fig. S2**). Then we tested SUI and MUI recordings that showed significant preference to a specific temporal position and confirmed that the information encoded by FR is preserved independently of the epoch length (**Fig. 2D)**. We did that by testing if the normalized FRs across temporal positions, when taking into account the whole period, is correlated with the normalized FRs when limiting analysis to the SED. The FRs correlation was strong and significant (r = 0.79, p < 4·e^-38^). Moreover, we confirmed that the preferred temporal position was the same for either length (**Fig. 2D, right panel)**. Even when applying this FR correlation analysis between the whole epoch duration and a shorter period of 4.5 seconds aligned with the chest opening (last 1.5 seconds are the object presentation), the preference was preserved (**Fig. 2E)**, with a significantly positive correlation (r = 0.74, p < 9·e^-31^). Thus, the FRs preference was preserved across the whole navigation and object collection period. Since the identity of objects changed on each trial, the FR preference exhibited by these neurons relates to their different temporal positions and not the objects’ identity, reflecting temporal order encoding at the single-neuron level in the human MTL.

Next, we analyzed the prevalence of neurons encoding temporal order in different regions of the MTL. Our results replicate the temporal selectivity reported recently in amygdala (AMY), entorhinal cortex (EC), and hippocampus (HC) (*9*, *10*), but also show that neurons in parahippocampal cortex (PHC) and temporal pole (TP) exhibited similar selectivity. We evaluated 294, 160, 218, 114 and 149 units from AMY, EC, HC, PHC and TP, respectively, which showed 44, 37, 45, 19 and 23 units that were modulated by temporal order (**Fig. 2Fi**). Among these units, there were 93, 65, 74, 38 and 42 SUIs recorded from AMY, EC, HC, PHC and TP, respectively, among which 18, 11, 18, 6 and 6 units were significantly modulated by temporal order. This corresponds to 19%, 17%, 24%, 16%, and 14% of the neurons being temporal order cells, a selectivity higher than chance level (p < 3·e^-07^, 1.6·e^-04^, 5.6·e^-09^, 0.005, and 0.009; one-sided binomial tests, **Fig. 2Fii, lower panel**). We recorded 201, 95, 144, 76 and 107 MUIs, from AMY, EC, HC, PHC and TP, respectively, which showed 26, 26, 27, 13 and 17 MUIs with selectivity to temporal order resulting in ratios of 13%, 27%, 19%, 17%, and 16%, which is significantly higher than 5%-selectivity produced by chance (p < 6·e^-06^, 1·e^-13^, 1·e^-09^, 4·e^-05^ and 1·e^-05^; one-sided binomial tests, respectively, **Fig 2Fii upper panel**). Both SUIs and MUIs showed a preference to be more selective to the first and third position (**Fig. 2Fiii, upper panel**), which may reflect primacy and recency effects at the single neuron level, as previously reported (*10*). When we tested whether FR is selective to one temporal positon against the rest after passing the surrogate test, post-hoc tests (FR of preferred temporal position > rest of temporal positions; Wilcoxon rank sum test, p < 0.05) showed that 53% and 48% of the modulated SUIs and MUIs preferred one temporal position over the rest. In total, these results suggest that the FRs of the temporal position selective population contain information about the temporal position of the object collected from treasure box in the TH task.

In a next step, we thus asked whether we would be able to decode the temporal position from the FRs of these neurons. To do so, we utilized a support vector machine (SVM) classifier that used the neurons’ FRs as features to classify the temporal positions. We constructed the classifier analysis in a binary way, classifying one position against the others. Thus we repeated this analysis three times: first position versus the rest, second versus the rest and third versus the rest. We trained the classifier on 80% of the data and tested on the remaining 20%. We repeated this procedure 10 times for the classification of each temporal position, using randomly selected 80% and 20% to cross-validate our results. Our results confirmed that the FRs can classify the temporal positions significantly better than the classification obtained after shuffling the labels of the temporal position (**Fig. S3A**) with an accuracy of 88.9% ± 9.6% (mean ± std), 70.7 ± 8.6%, and 78.6 ± 5.8%, with (p < 0.001, p = 0.11 and p = 0.036; permutation test) for first, second and third temporal position, respectively. The results for the area under the curve (AUC) were 96.4 ± 4.3 (mean ± std), 76.5 ± 6 and 80.6 ± 8, with (p = 0.001, p = 0.069 and p = 0.033; permutation test) for the first, second, and third temporal position, respectively. These results show better classification for the first and third temporal position, which presumably stems from the overrepresentation of these positions by individual neurons (**Fig. 2Fiii, upper panel**). This overrepresentation also affects the accuracy of the same SVM classification using FRs of MUIs which was 95.7±6% (mean ± std), 76.4 ± 4.8% and 82.1 ± 7.7%, with (p < 0.001, p = 0.048 and p = 0.006; permutation test) for the first, second, and third temporal position, respectively (for more details see **Fig. S4A**).

After establishing how temporal order is encoded by MTL neurons, we examined whether MTL neurons might also play a role in reading out the temporal order associated with the collected items. When the patients answered temporal retrieval questions, as shown in **Fig. 3A**, they were asked to recall which of two objects was presented later in the preceding encoding period. When the trial had three objects, this required three retrieval questions to cover all object combinations, while trials with two objects required just one question.

**Figure 3.**
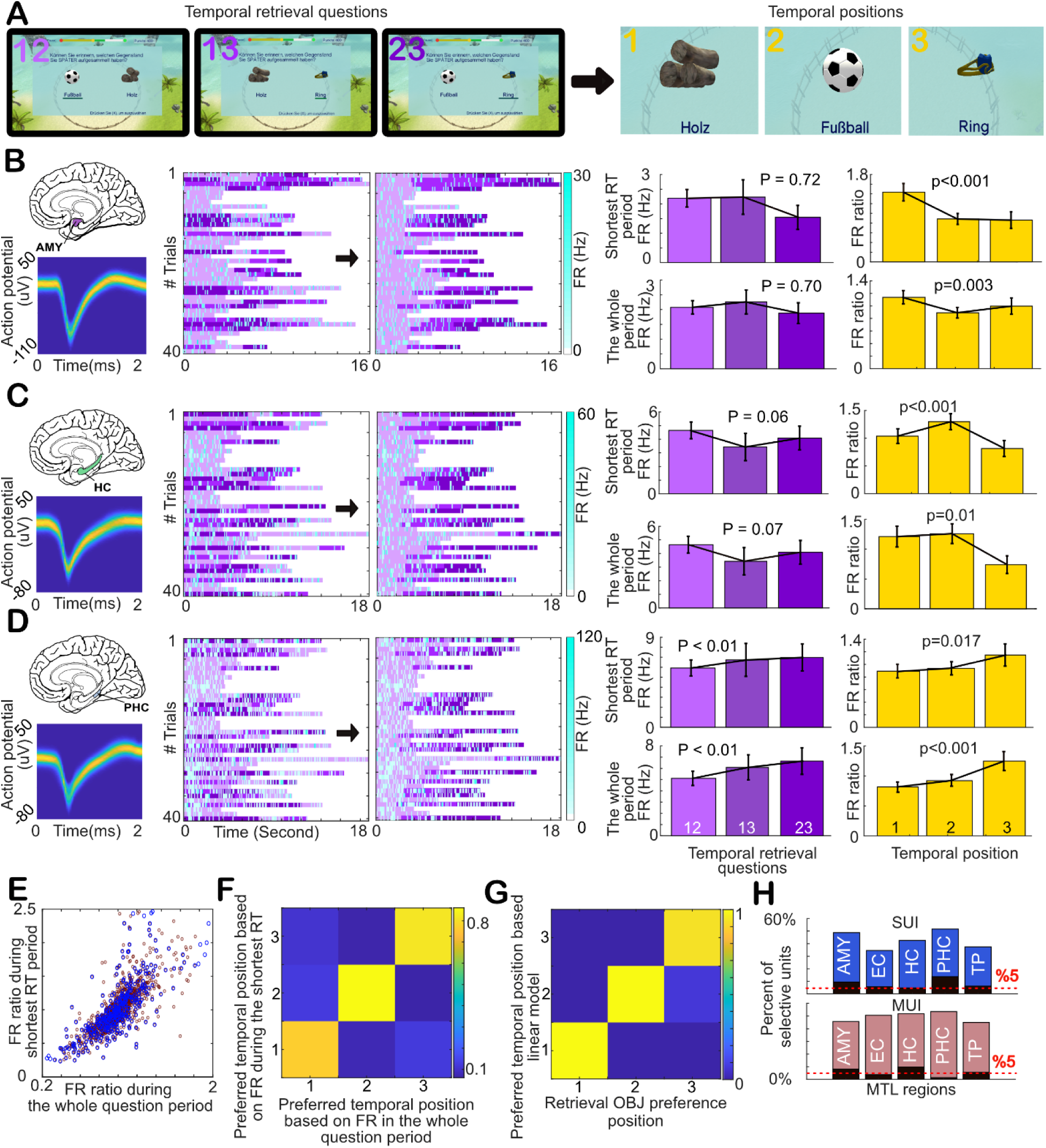
Neurons in the MTL are selective to temporal positions during retrieval. **A**. Illustrates a sequence of questions which ask the subject to recall which of two objects was presented later in a given trial. When the trial has three objects, it requires three retrieval questions as shown in A. Trials of two objects require just one question. **B**. (up) Illustrates the region in which the neuron was recorded at and (low) Plots the spike waveform as a density function. The next two panels to the right plot the spiking events of the neuron within 100ms bins (cyan), superimposed on the temporal retrieval questions in purple (very light, light to dark purple for questions 12, 13, and 23, respectively). The first panel shows the questions’ order in the experiment and the second panel shows them reordered (12, 13 and 23). The bar plots in purple illustrate the firing rate of the neuron during different questions, during the whole question period (low), or during the time required for the shortest reaction time (RT) (up). The bar plots in yellow show the FR ratio of when an object (associated with temporal position) is presented divided by the FR when it is not presented. Note that spiking events of the neuron in B are concentrated in q12 and q13, indicating that the neuron gets 1.5 times as active in questions containing objects associated with the first temporal position compared to questions that do not contain it. This FR ratio is significantly higher than FR ratios of objects presented in the second or third temporal position, which are not reaching 1 here. Note that the p values in the bar plots are based on surrogate statistics, in which the raw F-value is compared to the surrogate F-values, similar to Fig. 3. **C** and **D** present neurons that show a significant selectivity to the objects presented in the second and third temporal positions, respectively. **E**. Plots the normalized FR ratios for all temporal positions. The x-axis plots FR ratios calculated for the whole questions periods against y-axis of FRs ratios calculated for the time required for the shortest RT. Thus each neuron provides three data-points, one for each temporal position. Note the significant correlation between the FR ratios in both time windows. The highest FR ratio across temporal positions stays constant independently of the chosen time interval, as indicated by the heat plot in **F. G.** A heat plot showing the preferred temporal position, based on the FRs ratio during the shortest RT on the x-axis, against the preferred temporal position during retrieval, based on a mixed linear model for the same time period (see methods for more details). Note that the estimated preferred temporal positions are consistent across both methods. **H**. Illustrates the percentages of significantly selective MUI and SUI recordings from each region based on FRs ratio analysis in pink and blue respectively, and based on mixed linear model method in in black.

To directly evaluate the neurons’ engagement in temporal memory recall, we tested whether the FR of a given neuron increased significantly when an object associated with a specific temporal position was presented in a question as compared to not being presented. For example a neuron recorded in AMY (**Fig. 3B**) showed a relatively higher FR for questions that included objects associated with first temporal position with 2.1 ± 0.2 Hz (mean ± ci) and 2.3 ± 0.4 Hz for questions including objects of temporal positions 1 and 2 (“q12”) or 1 and 3 (“q13”), respectively, vs. 1.9 ± 0.4 Hz for questions including objects of temporal positions 2 and 3 (“q23”). These FR differences were not significant (p = 0.70; ANOVA-surrogate statistics). This was not surprising, given that we expected the neurons to be selective to objects associated with temporal position and not to the questions themselves which included two objects presented at different positions. So, to infer the FR associated with a specific temporal order, we took the FR during the questions that presented objects associated with a specific temporal position and divided this FR by the average FR during questions that did not include an object associated with that position. For example, to infer the FR associated with the first temporal position, we took the FRs during the q12 and q13 and divided it by the averaged FR during q23. We refer to the outcome as *FR ratio.* This procedure enabled us to see a significant preference to the first temporal positon with a ratio of 1.13 ± 0.11, 0.9 ± 0.1 and 1.0 ± 0.13 (mean ± ci) for the first, second and third position, respectively (p=0.005; ANOVA- surrogate statistics).

We then tested whether the entire question presentation period was needed to read out this information, or if the initial period of the question presentation already contained sufficient temporal information. We repeated our FR ratio analysis for the shortest reaction time (RT) that was needed to answer a question in a session (e.g., 1.3 seconds). Restricting the analysis to these 1.3 seconds showed that the early period of the question presentation already contained sufficient temporal information for classification, whereas the averaged RT the patient needed to answer a question was 3.9 ± 6.7 seconds (mean ± std). For the shortest RT, we observed similar FRs for the questions with 2.2 ± 0.3 Hz, 2.2 ± 0.6 Hz and 1.5 ± 0.4 Hz for q12, q13 and q23, respectively (same example neuron as above). The FR ratios similarly indicated a preference to the first temporal position with 1.4 ± 0.2, 0.9 ± 0.1 and 0.87 ± 0.17 for the first, second, and third position, respectively (p < 0.001; ANOVA-surrogate statistics). Other neurons preferred different temporal positions during recall. For the neurons in **Fig. 3C and D**, recorded from HC and PHC, we observed that their FR ratios for the shortest RT indicated their preference for the second and third temporal positions, respectively (p < 0.001 in both cases; ANOVA- surrogate statistics). When including the whole question periods the results remained the same (p < 0.001 in both cases; ANOVA-surrogate statistics). This selectivity was not only seen for SUIs but also for MUIs (**Fig. S4**).

When comparing the FR ratios for the whole question period with the FR ratios during the shortest RT, for both SUI and MUI, we found a strong and significant correlation (Pearson’s correlation; r = 0.76, p = 2·e^-238^; **Fig. 3E**). The preferred temporal positions also stayed the same for both periods (**Fig. 3F**). Thus, the information about temporal order appears to be available in the MTL before reporting the answer, potentially shortly after the visual perception of item identity. This suggests an association between each item’s identity and its temporal position. We compared the results of the FR ratio analysis with the result produced from linear mixed models (LMM) to infer the preferred position during recall. LMM fits the FR of a SUI or MUI to a model that have four linearly added components, three of which represent a particular temporal position as an independent variable, and the fourth component representing random effects (see methods). When comparing the preferred temporal position indicated by the FR ratio approach with the preferred temporal position indicated by LMM, we observed similar preferences. However, most of the neurons did not show a significant modulation in the LMM. Hence, when comparing the preferred positions resulting from both methods, we referred to the significant neurons using FR ratios independently of the significance level resulting from LMM analysis (**Fig. 3G**). We confirmed that our FR ratio analysis did not produce false-positive detections of modulated neurons by feeding the pipeline with shuffled question labels for all our recorded neurons. The detection rate was 9 neurons out of 312 with a percentage of 3%, which is similar to chance level (see methods).

All the MTL regions we included in our analysis showed a significant involvement in temporal position recall (41, 18, 26 15 and 18 out of 84, 52, 61, 29 and 48 SUIs) recorded from AMY, EC, HC, PHC and TP, respectively. The involvement of these regions exceeded 5%-chance level with percentages of 49%, 35%, 43%, 52% and 38%, respectively (p < 9·e^-32^, 5·e^-12^, 4·e^-19^, 1·e^-13^ and 1·e^-12^; one-sided binomial tests, respectively, **Fig. 3H**).

As for encoding, these results suggest that the relative FRs of MTL neurons reflect the temporal position of the collected item when the temporal recall question is answered. Hence, we tested whether we would be able to decode the temporal position from the relative FR ratios of MTL neurons. As for the encoding period, we utilized a SVM classifier that used relative FRs of the neurons showing temporal selectivity during recall as features to classify the temporal position during recall (same procedure as in the decoding analysis for the encoding period). Our results confirmed that the relative FRs can be used to classify the temporal position significantly better than the classification produced by shuffling the labels of the temporal position (**Fig. S3B**) with an accuracy of 91% ± 6.5%, 97.8% ± 2.3 96.1% ± 4.3% for the first, second, and third temporal positions respectively (p < 0.001 for all positions; permutation test). The AUC was as high with 0.987 ± 0.019, 0.994 ± 0.009, and 0.995 ± 0.0124, respectively, which is significantly higher than AUC produced by chance (p < 0.001 for all positions; permutation test). This classifier did extremely well, much better than the one used for encoding analysis, which may be due to the higher number of trials and selective neurons we found during recall than during encoding (110 selective neurons and at least 116 trials per neuron during recall, compared to 58 selective neurons and at least 70 trials per neuron during encoding. To control for data size differences, we repeated the SVM procedure with trial and neuron numbers matching those of the encoding period. We randomly selected 58 neurons and 70 trials, repeating the procedure 10 times. The results of this procedure (**Fig. S5**) showed decreases in the decoder accuracy and AUC, yet decoding of the second and third temporal positon during recall was still better than during encoding. The accuracy of the SVM classifier was 83.5 % ±4.1%, 91.4 % ±2.3% and 93.4% ± 2.3% for first, second, and third temporal positon, respectively, with significant difference for second and third temporal positions (p = 0.36, p = 0.00015, and p = 0.002; two-sample rank-sum test). The AUC was 0.9 ± 0.035, 0.98 ± 0.0179 and 0.98 ± 0.012, being significantly higher for the second and third temporal positions too (p = 0.18, p = 0.0001, and p = 0.026; two-sample rank-sum test). Running the same SVM classification using FRs of MUIs produced similar results to the ones in SUIs (**Fig. S4B**).

As with SUI recordings, MUIs recordings indicated similar MTL involvements in temporal memory recall (84, 40, 69 39 and 48 out of 268, 131, 194, 102 and 155 MUIs) recorded from AMY, EC, HC, PHC and TP, respectively. The involvement of these regions exceeded 5%-chance level with 46%, 51%, 52%, 53% and 45%, respectively (p < 2·e^-59^, 6·e^-32^, 4·e^-54^, 2·e^-32^ and 1·e^-34^; one-sided binomial tests, respectively). The LMM method showed the involvement of 15, 5, 11, 3 and 9 MUIs, respectively, with percentages of 8%, 6%, 8%, 4% and 8%, which were not higher than the chance level (p=0.05, p=0.41, p=0.07, p=1 and p=0.0840, respectively; one-sided binomial tests). When we tested whether FRs were selective to one temporal positon during recall against the rest after passing the surrogate test, post-hoc tests (FR of preferred temporal position > rest of temporal positions; Wilcoxon rank sum test <0.05) found 67% of modulated SUIs and 81% of modulated MUIs to prefer one temporal position over the rest during recall.

So far, we have described how MTL neurons contribute to encoding and recall of temporal positions. We next asked whether these two processes, encoding and recall, express a preserved temporal-order code at the cellular level. We hypothesized that it is more efficient to have a neuronal population using a preserved temporal order code in both processes. During encoding, the population must get activated by associating both inputs, the temporal position and the item associated with it. This is what we observe by the continuing activation during navigation across the whole epoch, including the presentation of the item (**Fig. 3D** and **E**). However, during recall, either input would trigger the activation of this population. This would allow the readout of the associated input. This is a likely scenario because both the encoding and recall neurons are distributed in overlapping MTL regions. However, in an alternative scenario, separate circuits may utilize different computational mechanisms. In such a scenario, no neurons are engaged in both processes, or even if they are engaged, they do not preserve the same preference for temporal positions. Here, we tested these scenarios by evaluating whether the neurons that were selective during encoding stayed selective during recall. Indeed, we observed a significant population of MTL neurons, represented in our SUI recordings, to preserve their selectivity in both processes. Two exemplary neurons (**Fig. 4A**), recorded from AMY and PHC, exhibited a preference to the first temporal position during both encoding and recall. The FR during the encoding was 1.7 ± 0.4 Hz, 1.1 ± 0.2 Hz, and 1 ± 0.5 Hz (mean ± ci) for the first, second and third position respectively, for the neuron 4A left with p < 0.001 ANOVA-surrogate statistics; p = 6 e^-9^ Wilcoxon rank sum test (first vs rest). The FR was 5.5 ± 0.5 Hz, 3.1 ± 0.4 Hz and 3.0 ± 0.6 Hz, respectively for the neuron 4A right with p < 0.001 surrogate statistics; p = 0.009 Wilcoxon rank sum test (first vs rest). During recall, the FR ratio was 1.29 ± .26, 0.97±0.22, and 0.86±0.22 (mean±ci) for the first, second and third position respectively, for the neuron 4A left, and 1.19 ± 0.13, 1 ± 0.13, and 0.87±0.13 for the neuron 4A right with p = 0.044 and p = 0.008 surrogate statistics for left and right neurons; p = 0.022 and p = 0.011 Wilcoxon rank sum test (first vs rest) for left and right neurons. Another two example neurons (**Fig. 4B**), recorded from EC and TP, showed a preference or tendency to prefer the second temporal position. During encoding, the FR was around (1.2 ± 0.3 Hz, 2.2 ± 0.3 Hz and 1.6 ± 0.5 Hz and 2.3 ± 0.4 Hz, 2.5 ± 0.3 Hz and 1.9 ± 0.5 Hz for the first, second and third position, for the neuron 4B left and right respectively, with p < 0.001 and p = 0.02 surrogate statistics; p = 0.22 and 0.15 Wilcoxon rank sum test (second vs rest) for left and right neurons respectively). During recall, the FR ratios were (0.99± 0.23, 1.21 ± 0.23, 0.88 ± 0.24, and 1.15 ± 0.2, 1.24 ± 0.2, and 0.78 ± 0.2 for the first, second and third position, for the neuron 4B left and right respectively with p = 0.15 surrogate statistics for left neuron and p = 0.014 ANOVA-surrogate statistics and 0.016 Wilcoxon rank sum test (second vs rest)) for the right. EC neuron in 4B showed a FR modulation tendency indicating a higher FR ratio to the second temporal position during recall, but this did not reach significance. Such neurons were pooled in an encoding pool that we analyzed afterward. Another two neurons (**Fig. 4C**) from PHC showed a preference or tendency to prefer the third temporal position. During the encoding, the FR was around (2.2 ± 0.3 Hz, 2.5 ± 0.5 Hz, and 3.4 ± 0.6 Hz, and 4.0 ± 0.8 Hz, 3.0 ± 0.4 Hz, and 4.4 ± 0.7 Hz for the first, second and third position, for the neuron **4C** left and right, respectively; with p<0.001 and p=0.01 surrogate statistics; p=0.002 and 0.027 Wilcoxon rank sum test (third vs rest) for left and right neurons respectively). In recall, the FR ratios were 0.89±0.11, 0.94±0.1, and 1.15±1.18, and 0.9±0.16, 0.9±0.14, and 1.18±0.24 for the first, second and third position, for the neuron **4C** left and right respectively; with p=0.02 and p=0.085 surrogate statistics for left and right neurons respectively; p=0.018 Wilcoxon rank sum test (third vs rest) for left neuron). The right neuron in 4C were pooled in encoding only too.

**Figure 4.**
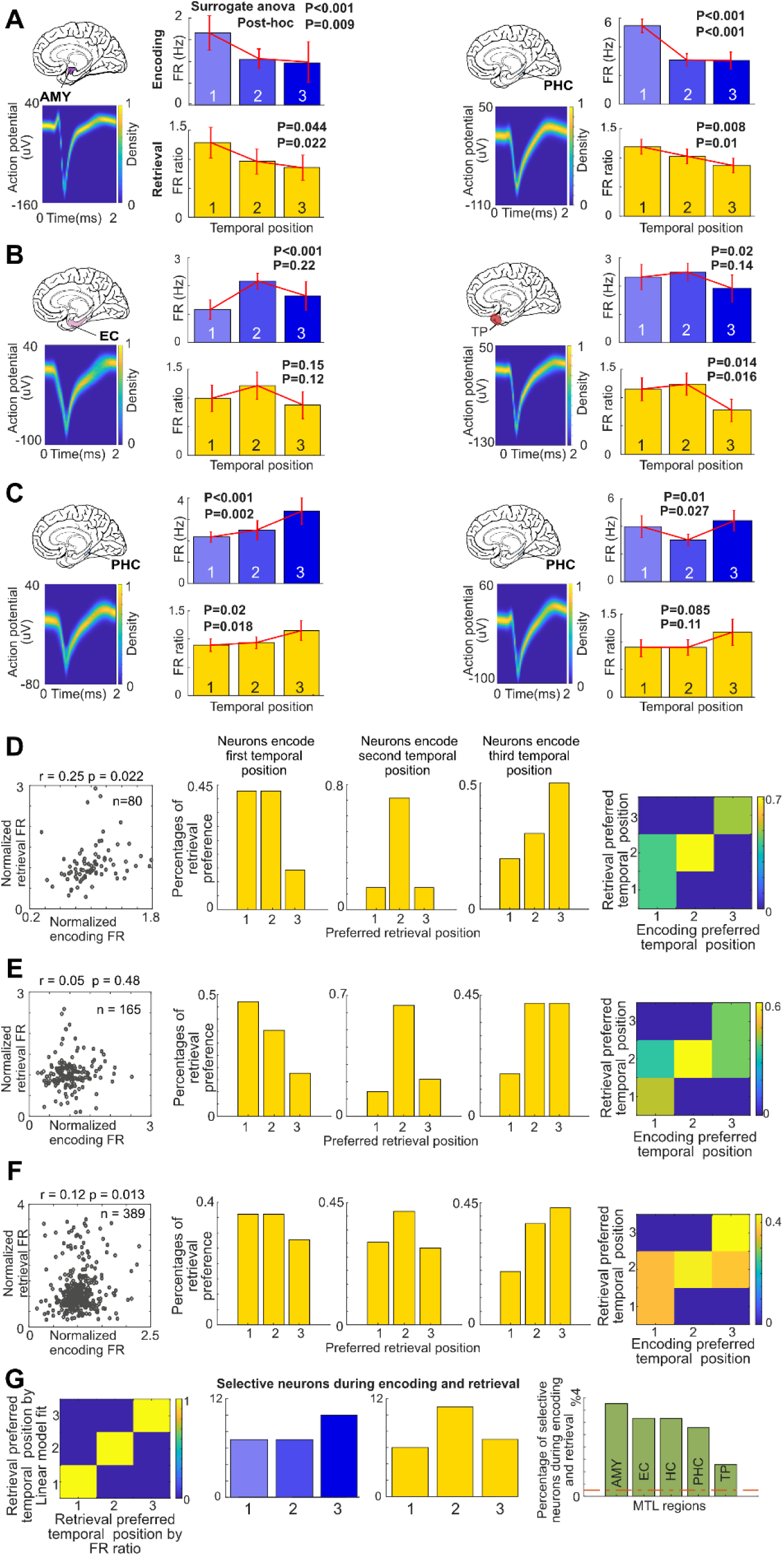
Neurons in the MTL which are selective to temporal position during encoding and retrieval preserve their selectivity. **A**. Presents two neurons that prefer the first temporal position during encoding and retrieval. The neuron in the left panel is recorded from AMY and the one in the right from PHC. Each side illustrates the region the neuron was recorded from (up) and plots the spike waveform as a density function (low). The bar plot in blue presents the neuron’s FR for each temporal position, averaged during the time required for the shortest epoch. The bar plot in yellow shows the FR ratios for each temporal position for the time required for shortest RT. The neurons presented in **B** and **C** illustrate preserved selectivity for the second and third position, respectively. **D**. Plots the normalized FR ratios against normalized FRs for the neurons which stayed significantly selective during encoding and retrieval. Note the significant correlation indicating a preserved selectivity to the same temporal position. The bar plots in yellow present the fraction of neurons with a preference for temporal positions during retrieval for neurons that were selective for first, second, and third temporal position, respectively during encoding. The bar plots indicate a trend to preserve the same temporal position. This is summarized in the heat map on the right that includes percentages exceeding the chance level of 33%. **E** and **F** replicate D but for the neurons which were selective during encoding, independent of retrieval, and F for selective neurons in retrieval, independent of encoding. Despite that both did not show correlation between FRs ratios and normalized FRs, both populations showed a tendency to preserve the selectivity to the same temporal position. **G**. Left is a heat plot the preferred temporal position during retrieval based on linear mixed model against the positions estimated by FRs ratios, for the neurons in D. The blue and yellow bar plots show the overall preferred temporal positions during encoding and retrieval for the neurons in D. The right plot shows the percentages of selective neurons during encoding and retrieval.

Hence, when we evaluated the whole population of cells that showed a significant modulation for different temporal positions (p<0.05; ANOVA-surrogate statistics, n=27) in both processes, we saw that the majority of these neurons preserved their preferred temporal position (**Fig. 4D**). This corresponds to the significant correlation between the normalized FRs across temporal positions during encoding with the FR ratios during recall (r=0.48, p=0.02; Pearson’s correlation). This correlation indicates that if a neuron had a high FR in one temporal position, it also exhibited a high FR when the item associated with that temporal position was presented in the recall question, providing support for our hypothesis of shared neural representations for temporal order between encoding and recall.

To further illustrate this result, we pooled these neurons based on their preference during encoding, and evaluated their preferences during recall (**Fig. 4D middle panels**). The yellow histograms indicate that the neurons preferring the first temporal position during encoding, also preferred it during recall, together with the second position. The neurons which preferred the second temporal position during encoding preferred the second during recall. Finally, the ones preferring the third position during encoding, also preferred the third position during recall. When summarizing these histograms in a confusion matrix plot for the bars that passed the chance level of 0.33. We see that the diagonal line is highlighted, demonstrating the preservation of the neuronal code of temporal positions (**Fig. 4D right panel**). We applied the same analysis on the neurons that showed significant modulation during encoding independent of recall. We did not see a significant correlation between the normalized FRs and the FR ratios (**Fig. 4E left panel**) for this population, but the histograms (**Fig. 4E middle panels**) and the confusion matrix (**Fig. 4E right panel**) still indicated that the temporal position preference was preserved to some extent. When testing the same procedure on the population which was significantly modulated during recall independently of the significance during encoding, we observed a significant correlation between the normalized FRs and the FR ratios (**Fig. 4F left panel**). We also found that the histograms and the confusion matrix (**Fig. 4E middle and right panels**) indicate that the temporal position preference is preserved, too, across encoding and recall. If we restricted our analysis to just the neurons that passed the post-hoc tests (FR of preferred temporal position > rest of temporal positions) after passing the surrogate test, we obtained similar results (**Fig. S7**). Thus, our results are not affected by the criteria of statistical testing. Moreover, when comparing the preferred retrieval temporal positions for the population in **D** with the preferred position result from LMM analysis, independent of the significant level of LMM, we see that the preferred positions are the same (**Fig. 4G left panel**). The overall preferred temporal positions for the population in **Fig. 4D** during encoding and recall (**Fig. 4G middle blue and yellow panels**) show a slight recency effect during encoding. The percentage of the neurons, which were significantly involved during both processes, encoding and recall, in MTL regions was significantly higher than chance level. We found 11, 5, 7, 3 and 2 selective neurons out of 293,158, 222, 108 and 155 neurons from AMY, EC, HC, PHC and TP, respectively. Thus, the resulting percentages were 3.8%, 3.2%, 3.2%, 2.8% and 1.3%, respectively, being significantly higher than the random chance level (0.05 for encoding multiplied by 0.05 for recall = 0.0025). All regions’ involvements exceeded random chance level (p = 4.1·e^-11^, 6.9·e^-06^, 2.4·e^-07^, 3.4·e^-04^ and 0.0143; one-sided binomial tests, **Fig. 4G right panel**).

Thus, we show here for the first time that, by preserving a selective firing to an item associated with a temporal position, MTL neurons may help to form and recall temporal order for a sequence of events by the same neuronal circuit. A straightforward computation that is ecologically efficient as it requires a low number of neurons to be involved. To proof that in practice, we trained the SVM classifier on all the encoding trials (n=70) using the FRs of the neurons (n=27) in **4C** as features. Using this for training, we successfully decoded the temporal positions associated with each item in the retrieval questions (n=112). The SVM classifier confirmed that the encoding FRs can be used to classify the temporal position during recall significantly better than the classification produced by shuffling the encoding FRs labels (**Fig. S3B**) with accuracy equals 60%, 56%, 66% for first, second and third temporal positions respectively, with (p=0.001, p=0.043 and p<0.001 for first, second and third positions respectively; permutation test).

The AUC was around 0.774, 0.56 and 0.65 respectively, which was significantly higher for first and third than the AUC produced by chance with (p<0.001, p=0.15 and p=0.003, respectively; permutation test). This is a very striking and important test because the classifier never saw the FRs of these neurons during recall except during testing, and because the recall trials (test trials) are around double the number (n=112) of the encoding trails (training trials, n=70). This shows that the FRs preferences are robustly preserved across both processes. We repeated this procedure but by training the classifier on the relative FRs (FR ratios) during recall, and then tested whether we can decode the temporal position during the encoding period based on the FRs of the same neurons in **4D**. Indeed that worked out very well and the classifier showed accuracy of 59%, 64% and 81% which was significantly higher for second and third than accuracy produced by chance with (p=0.22, p=0.02 and p<0.001, respectively). The AUC was 0.669, 0.682, and 0.839 which was significantly higher than the AUC produced from shuffled labels (p=0.011, p=0.007, and p<0.001).

We repeated the SVM analysis on MUIs using a similar number of units and trials. The performance of the resulting classifier was much worse with respect to accuracy and AUC in comparison to the classifier trained on SUI data. When training the classifier on the encoding trials to decode the temporal position during recall, the accuracy was 64%, 52% and 63% which was significantly higher just for first and third temporal positions than the accuracy obtained after shuffling the labels (p=0.001, p=0.26 and p=0.002). The AUC was 0.71, 0.45, and 0.48 which was significantly higher just for the first temporal position than AUC obtained after shuffling (p<0.001, p=0.8, and p=0.064). When flipping the process to training using the recall part to decode the temporal positions during encoding periods the accuracy was 54%, 60% and 49%, not significantly different from the accuracy obtained after shuffling (p=0.86, p=0.17, and p=1). AUC was 0.405, 0.490, and 0.293, also not significantly different from AUC produced by shuffling (p=0.91, p=0.54, and p=0.99). This means that it is not a rate code alone but as well the network needs to identify which neurons fire each spike to preserve the temporal order an observation speaks against anatomical clusters of neurons that prefer the same temporal positions. The results of AUC was 0.405, 0.490 and 0.293 which was not significantly different than AUC produced by shuffling (p=0.91, p=0.54 and p=0.99). In total, this indicate that MUIs contain higher noise they offer less information to preserve temporal order as to SUIs, confirmed by SVM classification results and their preferred temporal position across encoding and recall (**Fig. S8**).

## Discussion

In this study, we demonstrated that neurons in the MTL play a crucial role in supporting temporal order recall. This finding bridges the gap between the established role of MTL neurons in temporal order encoding (*9*, *10*) and their involvement in the recall process. Notably, we observed that a larger proportion of MTL neurons are selective for temporal order during recall than during encoding. SVM classifier analysis further confirmed this, showing higher accuracy in decoding temporal order during recall compared to encoding, even after controlling for the number of neurons and trials. Interestingly, most neurons selective during recall were not selective during encoding, suggesting a transfer of temporal order information across different neural populations in the MTL.

We identified a subset of neurons that maintained their selectivity across both encoding and recall phases, possibly serving as the core for preserving and transferring temporal order information. Training the SVM classifier on this small group of neurons during encoding was sufficient to accurately decode temporal positions during recall, and vice versa. This indicates that these neurons contain the essential information needed to track temporal order throughout the entire process, from navigation to recall. These findings show that firing rate information can encode and recall temporal order; however, whether neuronal spike timing relative to theta oscillations (*7*, *8*) supports this mechanism remains to be tested. Firing rate information and spike-field interactions might offer complementary perspectives on the same underlying computations. The fact that a small number of neurons can efficiently encode temporal order suggests this mechanism could be universal across different types of associative memory. Similar rapid encoding of new memories by linking associated items has been observed at the single-neuron level in the human MTL(*12*), aligning with our findings of single-neuron associations between temporal order and objects. Long-term associations between items, or between items and locations, have been linked to mechanisms like spike time-locking to local field potential oscillations and ripples activation(*13*, *14*). Further studies are needed to explore how these mechanisms interact with firing rate computations in the context of temporal memory.

MUIs showed similar temporal order selectivity during both encoding and recall as SUIs. This was confirmed by an SVM classifier trained solely on MUA, which successfully decoded temporal positions during both processes. However, MUA did not maintain its selectivity across both encoding and recall, likely due to different neurons being selective in each process. This suggests that neurons in the MUA group that are active during one process prefer different temporal positions. This finding indicates that selectivity for temporal positions is not based on anatomical clusters, as observed in sensory and motor modalities (*15–17*). It implies that preserving temporal order relies not on a simple rate code but on the network’s ability to identify which neurons fire at specific times. If neurons in the same MUI cluster preferred the same temporal positions, we would expect better classification performance in MUI-based SVM analysis. Thus, the firing rate code of neurons with a specific temporal order preference is sufficient to elegantly preserve temporal order across encoding and recall using only a small number of neurons

The sustained selectivity of temporal order neurons throughout the entire navigation and object collection phases during encoding, and across the full recall question period, suggests that these neurons differ from time cells. Time cells are selective to specific moments in a task, firing at precise time points (*11*, *18*, *19*). In contrast, temporal order neurons track order over extended periods, providing a continuous representation of temporal order. The sequential activation of time cells, which can track different task epochs in a low-dimensional space (*20*), suggests potential interactions between both systems. Additionally, the consistent observation of temporal order selectivity across five MTL regions suggests a universal temporal memory code in the MTL, though the exact underlying mechanisms remain to be explored.

## Supplementary figures

**Figure S1.**
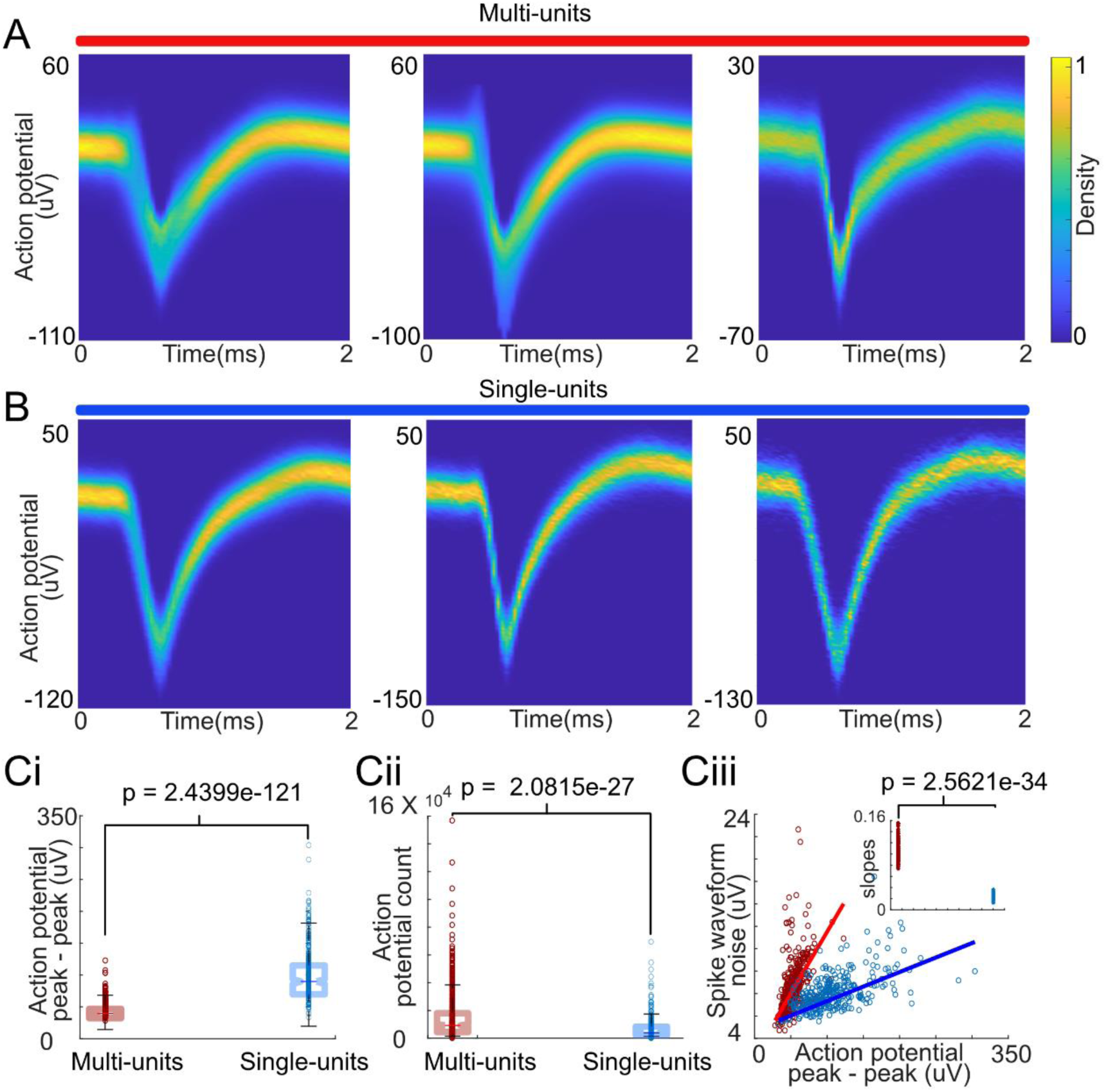
Evaluation of single- and multi-unit spike sorting. **A**. Plots density functions of three separate multiunits (MUIs). **B**. Plots density functions of three separate single units (SUIs). Note that the SUIs have a higher amplitude and a thinner waveform (i.e. less variability). **Ci**. Evaluates the peak to peak amplitude of the spike waveform of MUIs and SUIs in red and blue, respectively. **Cii**. Evaluates the action potential counts of MUIs and SUIs. Note that MUIs have higher numbers of counted action potentials. **Ciii**. Plots Spikes (action potentials) noise, measured by the standard deviation of the spike waveform, against the peak to peak amplitude. Note that the noise increases linearly with the amplitude but with much steeper slope for the MUIs than the SUIs. The small panel in Ciii plots 100 slopes estimated from a bootstrapping procedure that was repeated once for each slope to estimate it by fitting it to 100 randomly chosen data points from MUIs and SUIs. The 100 MUIs slopes are significantly steeper than the ones of SUIs indicating a noisier MUIs spike waveforms.

**Figure S2.**
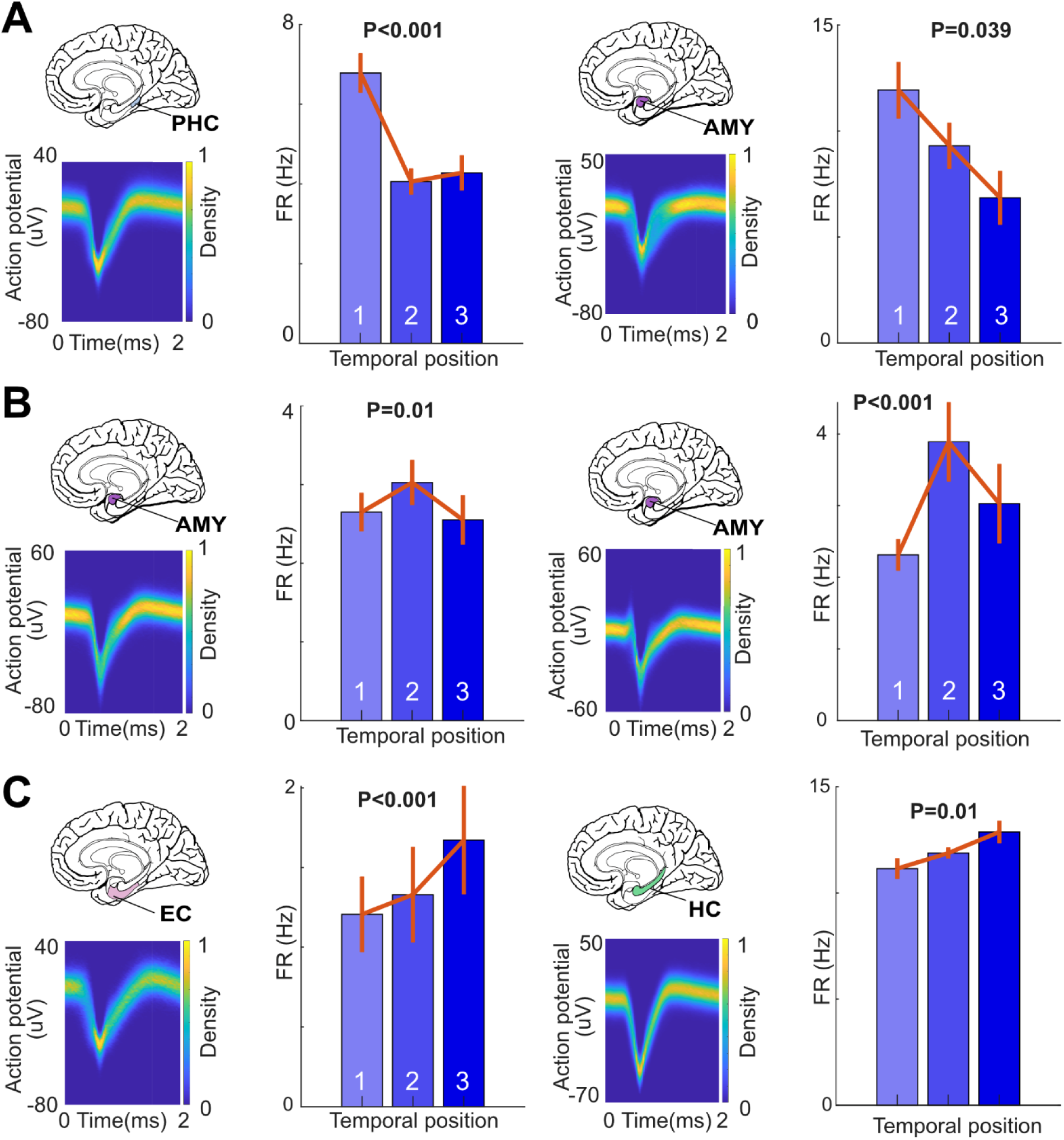
MUIs in the MTL are selective for different temporal positions during navigation. (**A, left**) Plots a density function of the spike waveform of a MUI recorded from PHC. The bar plot presents the averaged firing rate during each epoch taking the time needed for the shortest epoch, as in Fig. 2 (blue degree indicates the epochs 1, 2, and 3 using light, middle and dark blue respectively). The p-value indicates a significant modulation with higher FR for the first epoch (p<0.001; surrogate ANOVA). Note that the plots in A-right, show another MUI with similar selectivity recorded from AMY. The plots in B and C and repeating the same in A, but with a selectivity for the second and third temporal positions, respectively.

**Figure S3.**
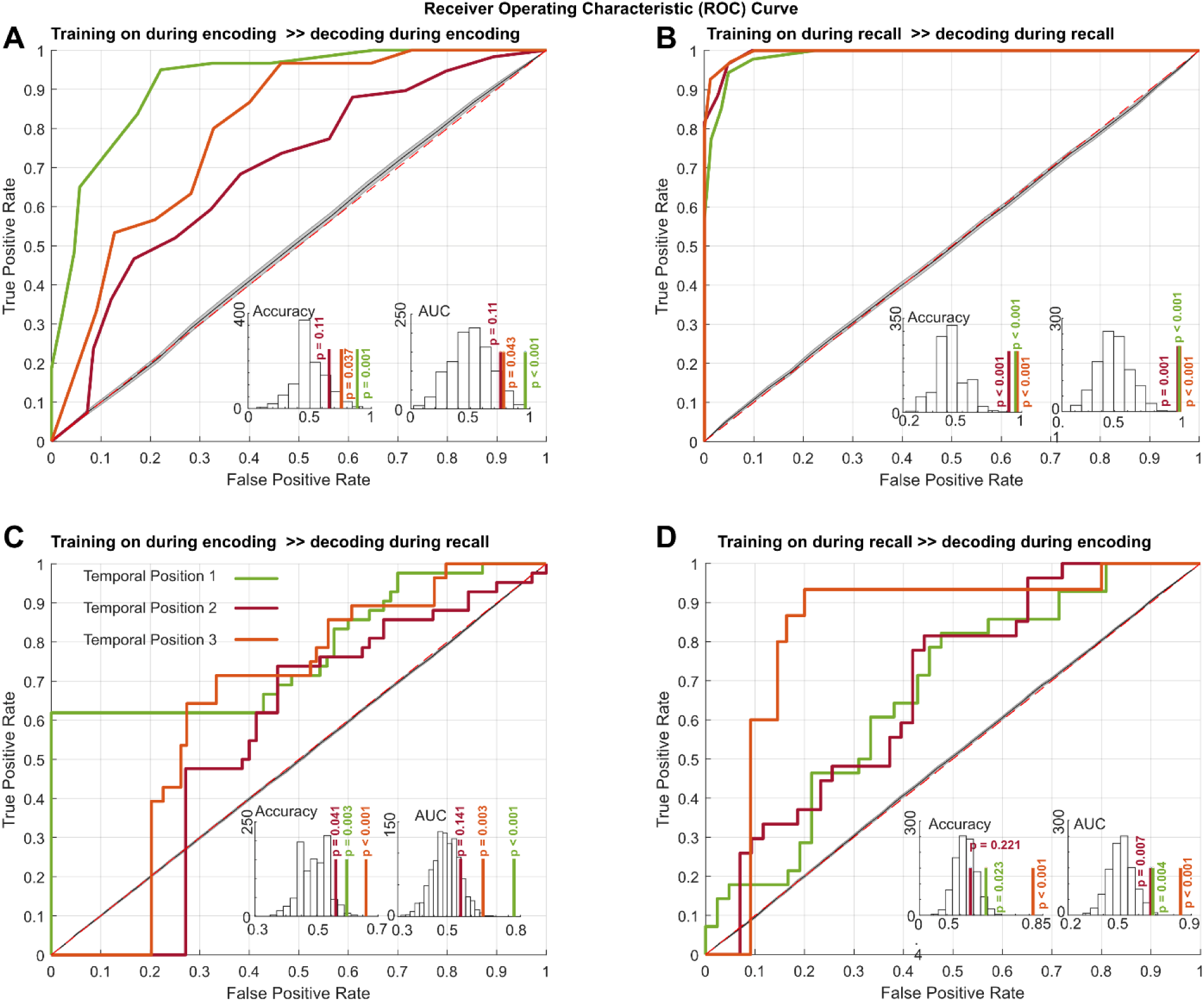
Classification results of a support vector machine classifier. **A.** Presents SVM classifier results for the encoding period, using FRs of selective neurons during encoding as features to predict temporal position. SVMs are used as binary classifiers; one temporal position against all the others, i.e., first position versus the rest (green), second versus the rest (red) and third versus the rest (orange). Cross-validation was performed by training the classifier on a random subset of 80% of the data and testing it on the remaining 20%, with 10 repetitions for each temporal position. The results confirmed that the FR features can be used to classify the temporal positions significantly better than by chance, estimated by the classification of shuffled labels with accuracy of 88.9 ± 9.6% (mean±std), 70.7±8.6% and 78.6±5.8%, with (p<0.001, p=0.11, and p=0.036; permutation test) for first, second and third temporal position respectively. The results for the area under the curve (AUC) was 0.964±0.043 (mean±std), 0.765±0.06 and 0.806±0.08, with (p=0.001, p=0.069, and p=0.033; permutation test) for first, second, and third temporal position, respectively. **B.** Similar to A, presents SVM classifier results for the recall period, using FR ratios of selective neurons during recall as features to predict temporal position during recall. The SVM classification accuracy of the temporal positions was significantly better than chance level, estimated by classification of shuffled labels with accuracy 91±6.5%, 97.8±2.3%, and 96.1±4.3% for the first, second and third temporal position respectively, with (p<0.001 for all positions; permutation test). The AUC was 0.987±0.019, 0.994±0.009, and 0.995±0.0124, respectively, which is significantly higher than AUC produced by chance with (p<0.001 for all positions; permutation test). Note that cross-validation was applied as in A. **C.** The SVM classifier was trained on the encoding periods of the trials, using the FRs of the selective neurons during encoding and recall as features. The classifier was tested on the recall period, aiming to decode the temporal positions. FR ratios of the same neurons were used as features for testing, i.e., features the classifier was not trained on. The SVM succeeded in classifying the temporal positions with higher accuracy than chance level estimated by classifying shuffled labels with accuracy 60%, 56%, and 66% for first, second and third temporal positions, respectively, with (p=0.001, p=0.043, and p<0.001 for first, second and third positions, respectively; permutation test). The AUC was around 0.774, 0.56, and 0.65, respectively, which was significantly higher for first and third temporal position than AUC produced by chance with (p<0.001, p=0.15 and p=0.003, respectively; permutation test). **D.** Represents the same procedure line as C, but the classifier used here was trained on FR ratios during recall and tested on FR during encoding. The classifier showed accuracy of 59%, 64%, and 81%, which was significantly higher for second and third temporal position than accuracy produced by chance with (p=0.22, p=0.02, and p<0.001, respectively). The AUC was 0.669, 0.682, and 0.839, which was significantly higher than the AUC produced after shuffling labels (P=0.011, P=0.007 and P<0.001).

**Figure S4.**
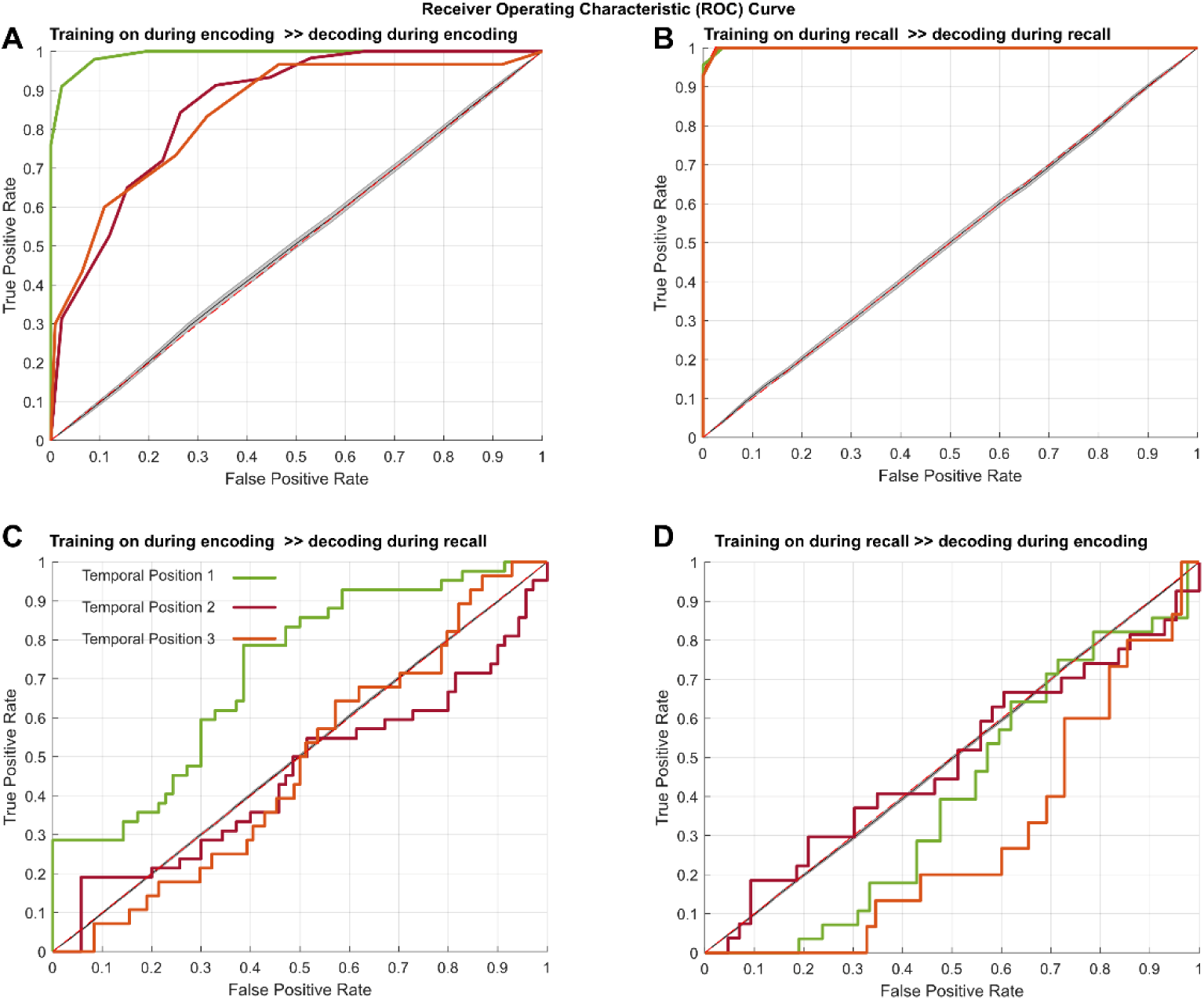
Classification results of a support vector machine classifier applied to MUIs. This figure repeats the same analysis as in S3, but with MUI data. **A** Accuracy of 95.7±6% (mean±std), 76.4±4.8% and 82.1±7.7%, with (p<0.001, p=0.048, and p=0.006; permutation test) for first, second and third temporal position, respectively. The results for the area under the curve (AUC) was 0.993±0.015 (mean±std), 0.857±0.069 and 0.842±0.15, with (p<0.001, p=0.07, and p=0.017; permutation test) for first, second and third temporal position, respectively. **B**. Accuracy was 99.0 ±1%, 98.1 ±2%, and 99.9 ±2%, with (p<0.001 for all; permutation test). The AUC was 0.995 ±0.054, 0.993 ±0.035 and 0.991 ±0.026 with (p<0.001 for all; permutation test). **C**. Accuracy was 64%, 52%, and 63% which was significantly higher just for first and third temporal positions than the accuracy obtained after shuffling the labels (p=0.001, p=0.26 and p=0.002). The AUC was 0.71, 0.45, and 0.48 which was significantly higher just for the first temporal position than AUC obtained after shuffling (p<0.001, p=0.8, and p=0.064). **D.** When flipping the process, i.e., using the recall part for training to decode the temporal positions during encoding periods, the accuracy was 54%, 60% and 49%, not significantly different from the accuracy obtained after shuffling (p=0.86, p=0.17, and p=1). AUC was 0.405, 0.490, and 0.293, also not significantly different from AUC produced after shuffling (p=0.91, p=0.54, and p=0.99).

**Figure S5.**
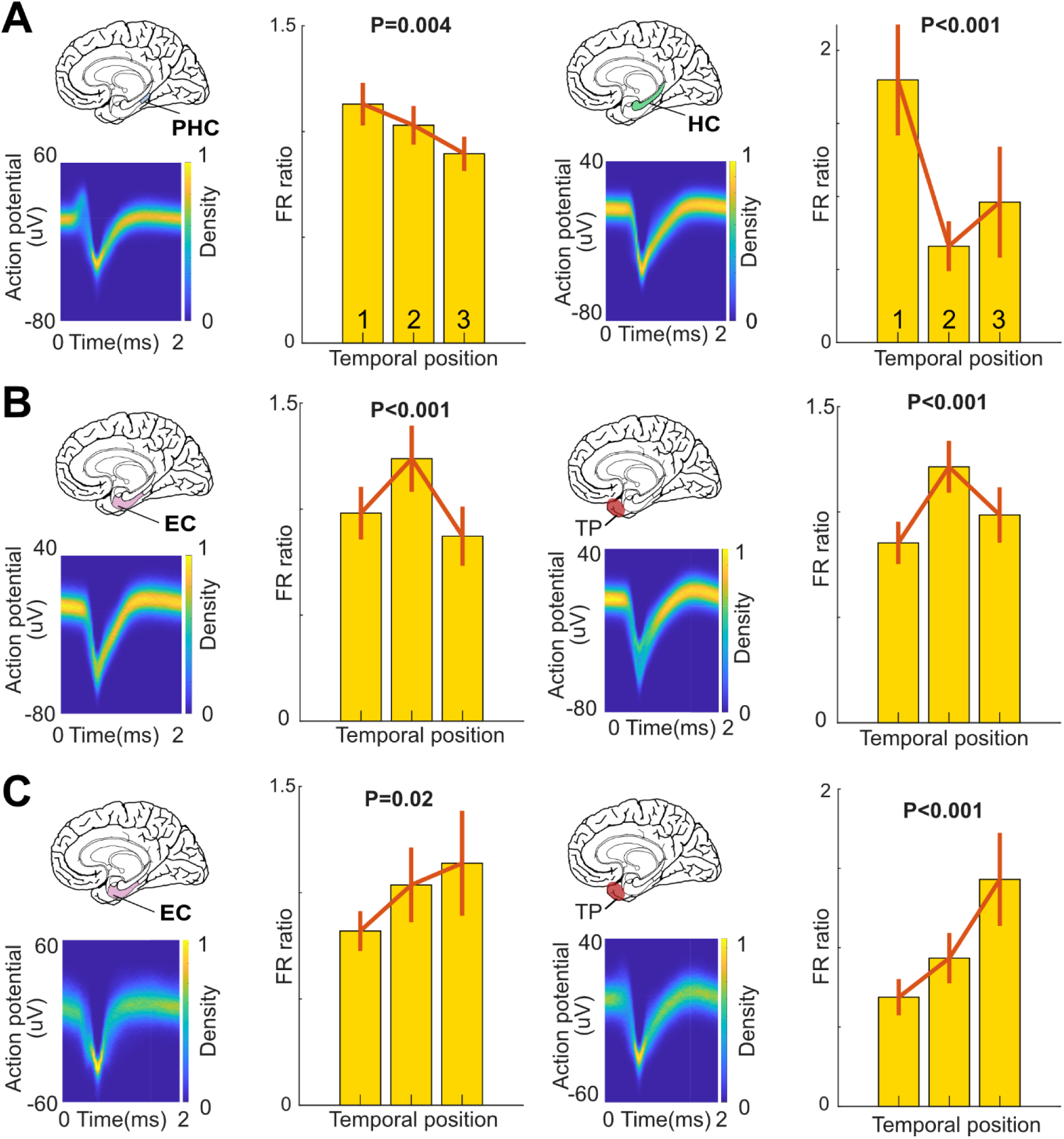
MUIs in the MTL are selective for different temporal positions during recall. This figure repeats the analysis shown in Fig. 3. **A-left**. Plots a MUI that is recorded from PHC. The density plot shows the spike waveform. The yellow bar plots in yellow show FR ratio of FR when an item (associated with temporal position) is presented divided by FR when that item is not presented. Note that this neuron is modulated towards higher FR when items associated with the first temporal position are presented in the question than items associated with the second or third position. Note that the p-values in the bar plots are based on surrogate statistics, in which the raw F value is compared to the surrogate F values, similar to Fig. 3. **A-right.** Presenting another MUI which is selective to the first temporal position. **C** and **B** replicate A with a significant selectivity for the items presented in the second and third temporal positions., respectively.

**Figure S6.**
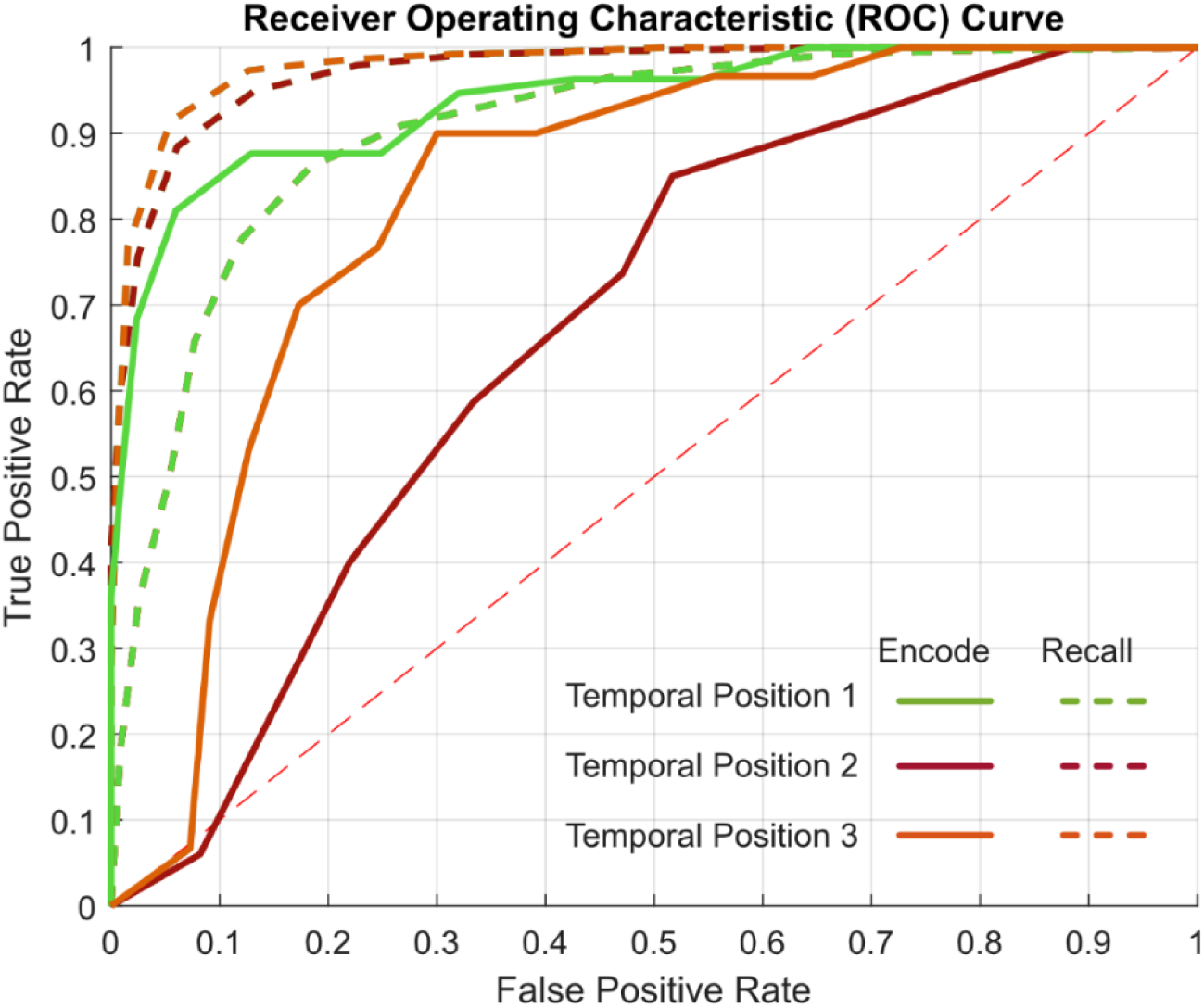
Comparison between the classification results during encoding and recall. This figure shows the SVM classification results for encoding (solid lines) and recall (disconnected lines) for the same data size. We randomly selected 58 neurons and 70 trials, repeated the classification 10 times, selecting 80% of the data set randomly for training, the remaining 20% for testing in each round. Note that SVM classification was better for recall than for encoding for the second and third temporal positions. The accuracy of the SVM classifier during recall was 83.5±4.1%, 91.4±2.3% and 93.4±2.3% for first, second and third temporal positon, respectively, after reducing the data size, with a significant difference for second and third temporal positions (p=0.36, p=0.00015, and p=0.002; two groups rank-sum test). The AUC during recall become 0.9±0.035, 0.98±0.0179, and p=0.98±0.012, significantly higher for second and third temporal positions, too (p=0.18, p=0.00018 and p=0.026; two groups rank-sum test).

**Figure S7.**
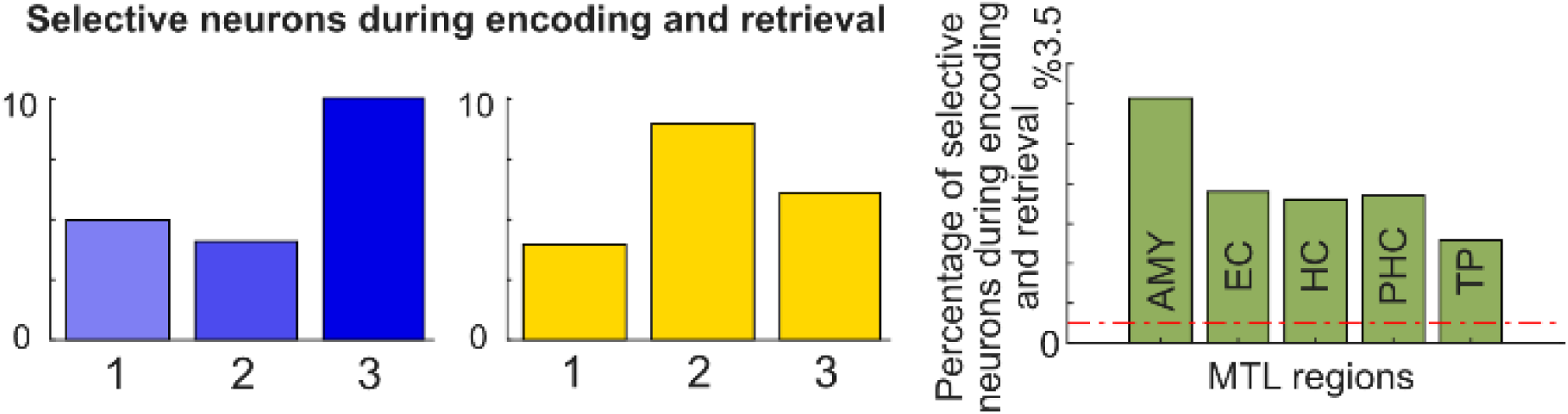
Quantification of neurons that are selective to one temporal position during encoding and recall. The blue and yellow bar plots show the overall preferred temporal positions during encoding and retrieval, respectively, for the neurons preferring one temporal position. The right plot shows the percentages of selective neurons during encoding and retrieval in all regions that have these neurons with a percentage higher than chance level (p<0.05; one-sided binomial tests).

**Figure S8.**
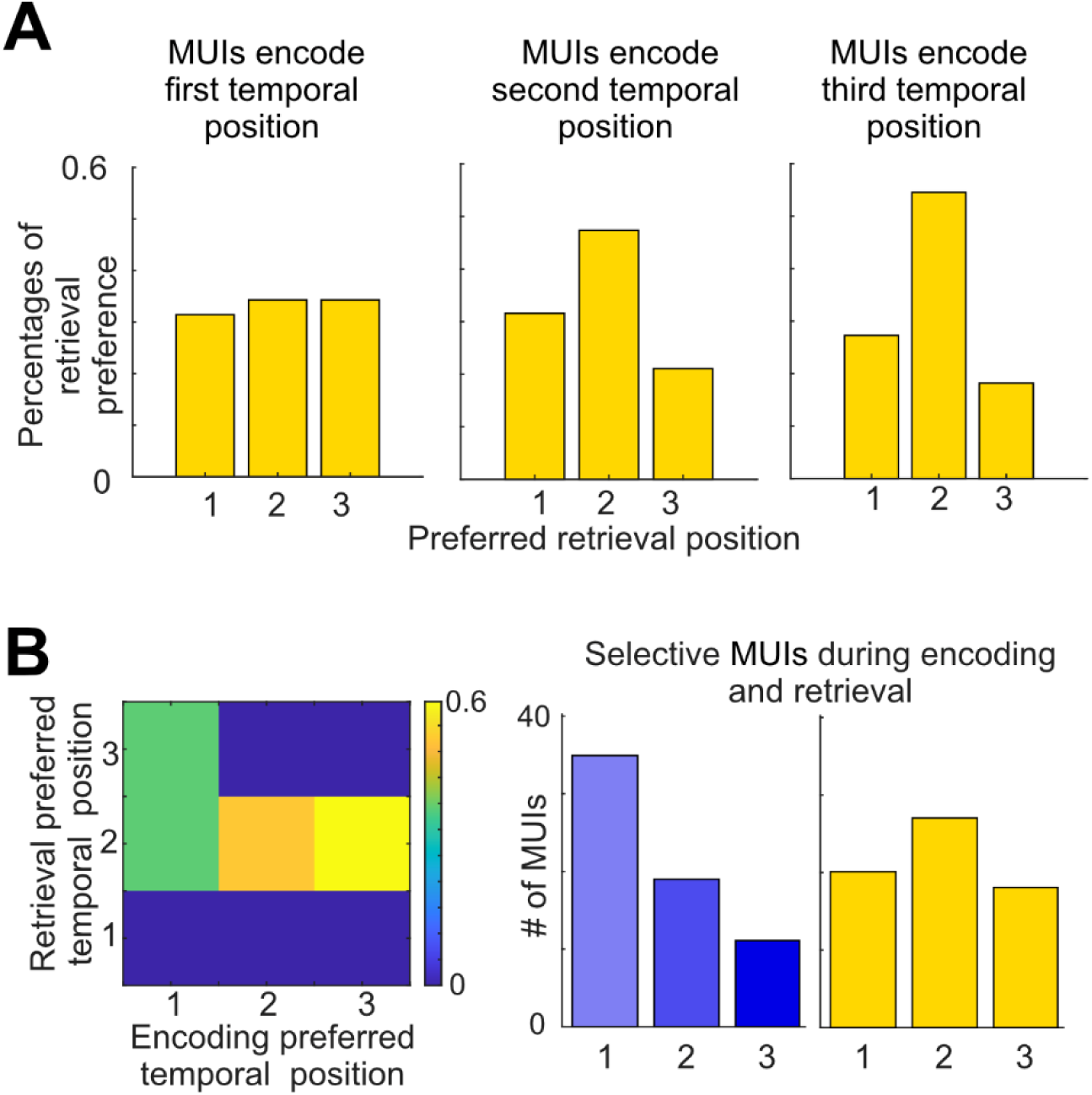
Quantification of MUIs that showed selectivity to temporal positions during encoding and recall. **A.** The bar plots in yellow present the percentage of MUIa that showed a preference for temporal positions during recall for neurons that were selective for first, second and third temporal position, respectively, during encoding. **B.** The heat map summarizes the bar plots in A including percentages exceeding the chance level of 33%. The blue and yellow bar plots show the overall preferred temporal positions for these MUIs during encoding and recall.

## Methods

### Human subjects

This study included data of 19 patients with pharmacologically intractable epilepsy who were implanted with depth electrodes at the Freiburg Epilepsy Center, Freiburg im Breisgau, Germany. Written consent conforming to the guidelines of the ethics committee of the University Hospital Freiburg, Freiburg im Breisgau, Germany was obtained from all patients.

### Neurophysiological

Patients underwent surgical implantation of intracranial depth electrodes in the medial temporal lobe to accurately identify the focus of epileptic seizures for potential resection. The specific locations and number of electrodes varied across individuals, based on clinical requirements. Neuronal signals were recorded using Behnke-Fried depth electrodes (AD-TECH Medical Instrument Corp., Racine, WI, USA), each consisting of a bundle of nine platinum-iridium microelectrodes with a diameter of 40 mm, extending from the tip of the depth electrode (*21*). Eight of the microelectrodes were used for electrophysiological recordings, which were later filtered to capture action potentials and local field potentials, while the ninth served as a reference electrode. Microwire data were recorded at a sampling rate of 30 kHz using the NeuroPort system (Blackrock Microsystems, Salt Lake City, UT, USA). The microelectrodes were implanted in various brain regions, including the amygdala, entorhinal cortex, fusiform gyrus, hippocampus, insula, parahippocampal cortex, temporal pole, and visual cortex.

### Spike detection and sorting

We used Wave_Clus (*22*) to detect and sort neuronal spikes. Each cluster was visually inspected and evaluated with respect to spike shape, its variability, and the presence of a clear refractory period based on inter-spike interval (ISI) distribution. We manually adjusted or excluded clusters where necessary. We excluded clusters with mean firing rates below 0.1 Hz during the analysis window (following (*23*, *24*)). We classified our clusters as single-unit when variability of their spike waveforms was low or multiunit for clusters with high waveform variability. This classification was quantified afterwards as shown in (**Fig. S1**). We identified 935 units, 623 multiunits and 312 single units. Neuronal responses from different sessions were treated as statistically independent. An experienced rater assigned depth electrode locations to brain regions using post-implantation MRI scans in order to link neurons recorded from each microelectrode bundle to a specific brain area.

### Spatial navigation–episodic memory task with embedded temporal memory

In the treasure hunt task, the patients sat in bed and performed a hybrid spatial navigation–episodic memory task running on a laptop. The task was adapted from previous studies (*9*, *25*) and implemented using Unity3D (Unity Technologies, San Francisco, CA, USA). The virtual environment was designed to resemble a beach, enclosed by a circular wooden fence with a diameter of 100 virtual units (vu). Navigation was restricted to the area inside the fence. There were no landmarks within the environment itself, but, some landmarks, such as palm trees and barrels, were positioned just outside the fence. One side of the beach bordered the sea, and the background featured mountains, palm trees, and the sky (Fig. 1A**-C**). Patients completed up to 40 trials per session. At the start of each trial, they were placed at a random location on the virtual beach (“passive home base transport”; Fig. 1A) and remained there until they initiated the trial by pressing a button. They then navigated to a series of treasure chests that appeared one after another on the beach (“navigation–encoding period”; Fig. 1A). Participants were encouraged to reach the chests as quickly as possible to earn bonus points for efficient navigation. Upon reaching a chest, they were automatically rotated to face it, and the chest opened to reveal an object and its name (Fig. 1A**-C**). After 1500 ms, the chest and object disappeared. Each trial involved visiting 2 or 3 chests, and across 40 trials, patients encountered 100 chests in total. After visiting the final chest in a trial, patients were passively transported to one of two elevated positions where they had an overhead view of the environment (“passive tower transport”; Fig. 1A**-C**). They then played a distractor game in which they tracked which of three moving boxes contained a coin. After the distractor game, the recall phase of the trial began. During the recall phase, patients either completed location-cued object recall or object-cued location recall on a trial by trial basis. We did not analyze these recall parts in this study. After recalling all locations and objects from a trial, patients completed a temporal order judgment task, where they were asked to decide which of two objects they had encountered later. Thus, the recall phase tested all components of episodic memory: object, location, and temporal information. At the end of the trial, patients received feedback on their performance, including points for correct object, location and temporal order recalls. Patients navigated the virtual environment using a game controller (forward, turn left, turn right), and their virtual positions and heading directions were recorded at 60 Hz. We synchronized the behavioral and electrophysiological data using triggers sent from the task paradigm to the recording system.

### General information on statistics

We used MATLAB 2020b with different MATLAB toolboxes and custom MATLAB code to analyze our data. The specifics of the statistical testing are indicated in the results, in general we considered results statistically significant when the corresponding p value fell below an alpha level of a = 0.05. Surrogate and permutation statistics were generally one-sided to assess whether an empirical test statistic exceeded a distribution of surrogate statistics significantly, unless otherwise specified. We adjusted the neuronal spike times to the behavioral time axis using the time stamps of trigger pulses that were sent from the laptop running the paradigm to the recording system. We then downsampled the behavioral data to 10 Hz following (*24*) and calculated the neuronal firing rate (Hz) for each sample (i.e., for each 100 ms time bin). We identified single- and multi-units to be temporally selective during encoding if they showed modulation of their firing rate across different epochs during navigation using one-way ANOVA with the factor temporal position. To measure how selective the units were during recall, we looked at their firing rates related to a specific object linked to a temporal positon. Since the recall question involved two objects at a time, we calculated the firing rate for a specific temporal position by checking firing rate of the units during questions with objects from that temporal position. Then, we divided this by the average firing rate during questions without objects from that temporal position. We named these values “FR ratios.” We calculated a FR ratio for objects associated with each temporal position. Then we compared the empirical F-value resulting from one-way ANOVA for a temporal position to 101 surrogate F-values resulting from the same ANOVA test of shuffled temporal position. The results of this test was checked using permutation testing in which the empirical difference between two FR ratios of two temporal positions was compared to 101 permutated difference value of the same positions after shuffling the labels. We applied this check three times per neuronal unit to cover the comparison between three temporal positions. We considered a unit selective if the empirical difference between any two temporal positions exceeded the 95th percentile of 101 permutations of the same positions. The difference between the permutation testing and the surrogate- ANOVA is that we apply one time ANOVA across all temporal positions to extract an F-value. In permutation testing, we extract the difference between every pair of temporal positions without any assumptions about the dependencies across the FR ratios of temporal positions. Each unit that was significant using the surrogate-ANOVA procedure was found significant in the permutation testing. In the linear mixed models (LMM) analysis, we inferred the preferred position during recall by LMM fits of the firing rates to a model that had four linearly added components, three representing each temporal position as effects, and one component representing random effects. We compared the empirical beta, being the maximum beta with surrogate beta generated by shuffling the temporal positions labels. We considered a unit significant if the empirical beta f exceeded the 95th percentile of 101surrogate beta factors. The preferred temporal position indicated by the FR ratio approach was similar to the preferred temporal position indicated by LMM (Fig. 3G and Fig. 4G), yet LMM showed lower percentages of significant units. The LMM analysis showed the involvement of (8, 3, 3, 4 and 3 SUIs, respectively) with percentage of 10%, 6%, 5%, 14% an 6%, that are higher than chance level for AMY and PHC only (p=0.049, p=0.52, p=0.73, p=0.03 and p=0.44; one-sided binomial tests, respectively). To test whether the surrogate-ANOVA or the permutation are causing false positives, we applied an additional control analysis. We shuffled the question labels for each unit and then fed it to our surrogate-ANOVA analysis pipeline. Since the data was shuffled across the questions, our pipeline should not detect significant neurons higher than a chance level of 5%. Indeed that was the case as we detected rate was 9 neurons out of 312, i.e. 3%, lower than chance level. We tested the stability of our analysis to the duration of epochs or the questions. We compared the normalized firing rates between different temporal positon for different duration, the start, the whole, or end of the periods. The selectivity stayed stable for both encoding and recall for the different periods (Fig. 2D).

### SVM decoding analysis

We employed support a vector machine decoder implemented in Matlab to test whether we can decode the temporal position from the firing rates of the selective single- and multi-units. We constructed the classifier analysis in a binary way, classifying one position against the others. We repeated this analysis three times: first position versus the rest, second versus the rest and third versus the rest. We trained the classifier on 80% of the data and tested on 20%. We repeated this procedure 10 times for each temporal position classification, using randomly selected 80% and 20% to cross-validate our results. Our results confirmed that the FRs can classify the temporal positions significantly better than the classification produced by shuffling the labels of the temporal position (**Fig. S3A**) with accuracy and area under the curve higher than chance level. Similar decoding was applied to data recorded during recall but with the FR ratios as predictors to decode the temporal positions. The amount of data used with the classifier during encoding is smaller than the amount of data collected during recall because more neurons showed stronger selectivity during recalls, and we collected more data-points per trial, as each trial had two to three recall questions. Therefore, the classifier had a much higher accuracy when decoding the temporal position during recall.

## Acknowledgments

We are grateful for the patients who participated in this study. We thank the clinical team of the Freiburg Epilepsy Center (Germany) for their continuous support. L.K., A.B., and A.S.-B. were supported by the Federal Ministry of Education and Research (BMBF; 01GQ1705A). J.J. received funding via National Science Foundation (NSF) grant BCS-1724243. L.K., A.B., and A.S.-B. were supported by NIH/NINDS grant U01 NS113198-01. L.K. received funding from the German Research Foundation (DFG; project no. 447634521) and was supported by a travel grant from the Boehringer Ingelheim Fonds (Mainz, Germany). P.C.R. received research grants from the Fraunhofer Society (Munich, Germany) and the Else Kröner-Fresenius Foundation (Bad Homburg, Germany). M.K. was supported by NIH grants MH55687 and MH061975. J.J. was supported by NIH grant MH104606.

## Author contributions

M.F.K and L.K. designed the study; L.K., A.B., P.C.R., and A.S.-B. recruited participants; P.C.R. implanted electrodes; L.K. and A.B. collected data; M.F.K. analyzed data with input from L.K., A.B., M.K, J.J., and A.S.-B. A.S.-B. secured funding and supervised the project; M.F.K. wrote the paper, and all authors reviewed the manuscript.

